# Consortia of anti-nematode fungi and bacteria in the rhizosphere of soybean plants attacked by root-knot nematodes

**DOI:** 10.1101/332403

**Authors:** Hirokazu Toju, Yu Tanaka

## Abstract

Cyst and root-knot nematodes are major risk factors of agroecosystem management, often causing devastating impacts on crop production. The use of microbes that parasitize or prey on nematodes has been considered as a promising approach for suppressing phytopathogenic nematode populations. However, as effects and persistence of those biological control agents often vary substantially depending on regions, soil characteristics, and agricultural practices, more insights into microbial community processes are required to develop reproducible control of nematode populations. By performing high-throughput sequencing profiling of bacteria and fungi, we examined how root and soil microbiomes differ between benign and nematode-infected plant individuals in a soybean field in Japan. Results indicated that various taxonomic groups of bacteria and fungi occurred preferentially on the soybean individuals infected by root-knot nematodes. Based on a network analysis of potential microbe–microbe associations, we further found that several fungal taxa potentially preying on nematodes [*Dactylellina* (Orbiliales), *Rhizophydium* (Rhizophydiales), *Clonostachys* (Hypocreales), *Pochonia* (Hypocreales), and *Purpureocillium* (Hypocreales)] co-occurred in the soybean rhizosphere at a small spatial scale. Overall, this study suggests how “consortia” of anti-nematode microbes can derive from indigenous (resident) microbiomes, thereby providing basic information for managing anti-nematode microbial communities in agroecosystems.

## INTRODUCTION

Plant pathogenic nematodes, such as cyst and root-knot nematodes, are major threats to crop production worldwide (Barker & Koenning 1998; Abad et al. 2008). Soybean fields, in particular, are often damaged by such phytopathogenic nematodes, resulting in substantial yield loss (Wrather et al. 1997; Wrather & Koenning 2006). A number of chemical nematicides and biological control agents (e.g., nematophagous fungi in the genera *Purpureocillium* and *Clonostachys*) have been used to suppress nematode populations in farmlands (Schmitt et al. 1983; Li et al. 2015). However, once cyst and root-knot nematodes appear in a farmland, they often persist in the soil for a long time (Meyer & Roberts 2002), causing high financial costs in agricultural management. Therefore, finding ways to suppress pathogenic nematode populations in agroecosystems is a key to reducing risk and management costs in production of soybean and other crop plants.

To reduce damage by cyst and root-knot nematodes, a number of studies have evaluated effects of crop varieties/species, crop rotations, fertilizer inputs, and tillage intensity on nematode density in farmland soil (Nusbaum & Ferris 1973; Thomas 1978; Barker & Koenning 1998; Okada & Harada 2007). However, the results of those studies varied considerably depending on regions, soil characteristics, and complicated interactions among multiple factors (e.g., interactions between organic matter inputs and tillage frequency) (Donald et al. 2009). Therefore, it remains an important challenge to understand the mechanisms by which phytopathogenic nematode populations are suppressed in some farmland soils but not others (Hamid et al. 2017). New lines of information are required for building general schemes for making agroecosystems robust to the emergence of pest nematodes.

Based on the technological advances in high-throughput DNA sequencing, more and more studies have examined structures of microbial communities (microbiomes) in order to evaluate biotic environmental conditions in the endosphere and rhizosphere of plants (Lundberg et al. 2012; Edwards et al. 2015; Schlaeppi & Bulgarelli 2015; Toju et al. 2018). Indeed, recent studies have uncovered microbiome compositions of “disease suppressive soils”, in which pests and pathogens damaging crop plants have been suppressed for long periods of time (Mendes et al. 2011; Berendsen et al. 2012; Mendes et al. 2013). Some studies have further discussed how some microbes within such disease-suppressive microbiomes contribute to health and growth of crop plant species (Mendes et al. 2011; Cha et al. 2016). In one of the studies, soil microbiome compositions were compared among soybean fields that differed in the density of cyst nematodes (Hamid et al. 2017). The study then revealed that bacteria and fungi potentially having negative impacts on nematode populations (e.g., *Purpureocillium* and *Pochonia*) were more abundant in long-term than in short-term monoculture fields of soybeans (Hamid et al. 2017). While such among-farmland comparisons have provided invaluable insights into ecosystem functions of indigenous (native) microbiomes, the farmlands compared in those studies could vary in climatic and edaphic factors, obscuring potential relationship between cropping system management and community processes of anti-nematode microbes. Moreover, because incidence of cyst and root-knot nematodes generally varies at small spatial scales (Gavassoni et al. 2001), there can be spatial heterogeneity in abundance and community compositions of anti-nematode bacteria and fungi within a farmland. Thus, studies focusing on fine-scale assembly of anti-nematode microbes are awaited for developing agroecosystem management protocols for controlling phytopathogenic nematodes.

By an Illumina sequencing analysis of bacteria and fungi in a soybean (*Glycine max*) field, we examined how root and rhizosphere microbiome structures varied among host plant individuals that differed in damage by root-knot nematodes (*Meloidogyne* sp.). Based on the data of microbiomes at a small spatial scale, we statistically explored microbial species/taxa that occurred preferentially in the roots or rhizosphere soil of nematode-infected soybean individuals. We further investigated the structure of networks depicting co-abundance patterns of microbial species/taxa within the soybean field, thereby examining whether multiple anti-nematode bacteria and fungi form consortia (assemblages) on/around the plant individuals infected by root-knot nematodes. Overall, this study suggests that various taxonomic groups of anti-nematode bacteria and fungi are present within indigenous microbiomes. Our results also suggest that microbiome assembly at fine spatial scales is a key to manage populations and communities of such functional microbes.

## MATERIALS AND METHODS

### Sampling

Fieldwork was conducted at the soybean field on the Hokubu Campus of Kyoto University, Japan (35.033 °N, 135.784 °E). In the field, the soybean strain “Sachiyutaka” was sown at 15 cm intervals in two lines (Supplementary Fig. 1) on July 4, 2016 [basal fertilizer, N:P_2_O_5_:K_2_O= 3:10:10 g/m^2^]. In the lines, 69 and 62 individuals (“set 1” and “set 2”, respectively), respectively, were sampled every other positions (i.e., 30 cm intervals) (Fig. 1) on October 7, 2016. The sampled soybean individuals were classified into three categories: normal individuals with green leaves (“green”), individuals with yellow leaves (“yellow”), and those with no leaves (“no leaf”) (Fig. 1A-C). Among them, “green” individuals exhibited normal growth, while “no leaf” individuals were heavily infected by root-knot nematodes: “yellow” individuals showed intermediate characters. In total, 97 “green”, 19 “yellow”, and 15 “no leaf” individuals were sampled (Fig. 1D).

**FIGURE 1.**
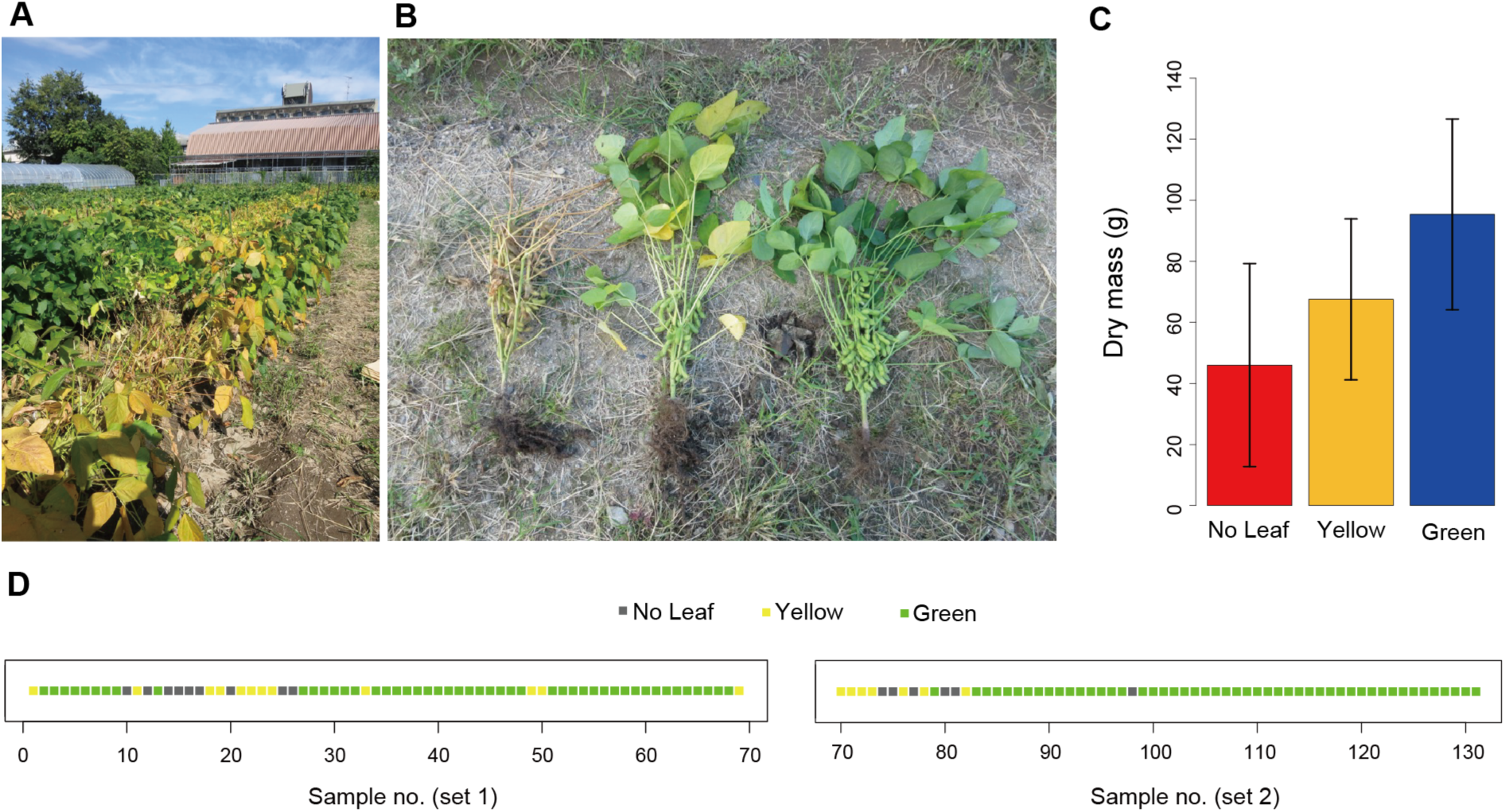
Study site and soybeans. **(A)** Soybean field in which sampling was conducted. **(B)** Soybean states. Soybean individuals were classified into three categories: those heavily attacked by root-knot nematodes (“no leaf”; left), those exhibited normal growth (“green”; right), and those showing intermediate characters (“yellow”; middle). **(C)** Relationship between soybean states and biomass. Dry mass significantly differed among “no leaf”, “yellow”, and “green” soybean individuals (ANOVA; *F*_2_ = 20.5, *P* < 00001). **(D)** Spatial distribution of “no leaf”, “yellow”, and “green” soybean individuals. Sampling sets 1 and 2 are shown separately.

For each individual, two segments of 5-cm terminal roots and rhizosphere soil were collected from ca. 10-cm below the soil surface. The root and soil samples were transferred into a cool box in the field and then stored at −80°C until DNA extraction in the laboratory. The whole bodies of the individuals were placed in drying ovens at 80 °C for 72 hours to measure dry mass. The dry mass data indicated that “green”, “yellow”, and “no leaf” soybean individuals significantly differed in their biomass (Fig. 1C).

### DNA Extraction, PCR, and Sequencing

The root segments of each individual were transferred to a 15 mL tube and washed in 70% ethanol by vortexing for 10 s. The samples were then transferred to a new 15 mL tube and then washed again in 70% ethanol by sonication (42 Hz) for 5 min. After an additional sonication wash in a new tube, one of the two root segments were dried and placed in a 1.2 mL tube for each soybean individual. DNA extraction was then performed with a cetyltrimethylammonium bromide (CTAB) method (Sato & Murakami 2008) after pulverizing the roots with 4 mm zirconium balls at 25 Hz for 3 min using a TissueLyser II (Qiagen).

For DNA extraction from the rhizosphere soil, the ISOIL for Beads Beating kit (Nippon Gene) was used as instructed by the manufacturer. For each sample, 0.5 g of soil was placed into a 2 mL microtubes of the ISOIL kit. To increase the yield of DNA, 10 mg of skim milk powder (Wako, 198-10605) was added to each sample (Takada-Hoshino & Matsumoto 2004).

For each of the root and soil samples, the 16S rRNA V4 region of the prokaryotes and the internal transcribed spacer 1 (ITS1) region of fungi were amplified. The PCR of the 16S rRNA region was performed with the forward primer 515f (Caporaso et al. 2011) fused with 3–6-mer Ns for improved Illumina sequencing quality (Lundberg et al. 2013) and the forward Illumina sequencing primer (5’-TCG TCG GCA GCG TCA GAT GTG TAT AAG AGA CAG-[3–6-mer Ns] – [515f] −3’) and the reverse primer 806rB (Apprill et al. 2015) fused with 3–6-mer Ns and the reverse sequencing primer (5’-GTC TCG TGG GCT CGG AGA TGT GTA TAA GAG ACA G [3–6-mer Ns] -[806rB] −3’) (0.2 μM each). To prevent the amplification of mitochondrial and chloroplast 16S rRNA sequences, specific peptide nucleic acids [mPNA and pPNA; Lundberg et al. (2013)] (0.25 μM each) were added to the reaction mix of KOD FX Neo (Toyobo). The temperature profile of the PCR was 94 °C for 2 min, followed by 35 cycles at 98 °C for 10 s, 78 °C for 10 s, 50 °C for 30 s, 68 °C for 50 s, and a final extension at 68 °C for 5 min. To prevent generation of chimeric sequences, the ramp rate through the thermal cycles was set to 1 °C/sec (Stevens et al. 2013). Illumina sequencing adaptors were then added to respective samples in the supplemental PCR using the forward fusion primers consisting of the P5 Illumina adaptor, 8-mer indexes for sample identification (Hamady et al. 2008) and a partial sequence of the sequencing primer (5’-AAT GAT ACG GCG ACC ACC GAG ATC TAC AC - [8-mer index] - TCG TCG GCA GCG TC −3’) and the reverse fusion primers consisting of the P7 adaptor, 8-mer indexes, and a partial sequence of the sequencing primer (5’- CAA GCA GAA GAC GGC ATA CGA GAT - [8-mer index] -GTC TCG TGG GCT CGG −3’). KOD FX Neo was used with a temperature profile of 94 °C for 2 min, followed by 8 cycles at 98 °C for 10 s, 55 °C for 30 s, 68 °C for 50 s (ramp rate = 1 °C/s), and a final extension at 68 °C for 5 min. The PCR amplicons of the 131 soybean individuals were then pooled after a purification/equalization process with the AMPureXP Kit (Beckman Coulter). Primer dimers, which were shorter than 200 bp, were removed from the pooled library by supplemental purification with AMpureXP: the ratio of AMPureXP reagent to the pooled library was set to 0.6 (v/v) in this process.

The PCR of fungal ITS1 region was performed with the forward primer ITS1F-KYO1 (Toju et al. 2012) fused with 3–6-mer Ns for improved Illumina sequencing quality (Lundberg et al. 2013) and the forward Illumina sequencing primer (5’-TCG TCG GCA GCG TCA GAT GTG TAT AAG AGA CAG-[3–6-mer Ns] – [ITS1-KYO2] −3’) and the reverse primer ITS2-KYO2 (Toju et al. 2012) fused with 3–6-mer Ns and the reverse sequencing primer (5’-GTC TCG TGG GCT CGG AGA TGT GTA TAA GAG ACA G [3– 6-mer Ns] - [ITS2-KYO2] −3’). The buffer and polymerase system of KOD FX Neo was used with a temperature profile of 94 °C for 2 min, followed by 35 cycles at 98 °C for 10 s, 50 °C for 30 s, 68 °C for 50 s, and a final extension at 68 °C for 5 min. Illumina sequencing adaptors and 8-mer index sequences were then added in the second PCR as described above. The amplicons were purified and pooled as described above.

The sequencing libraries of the prokaryote 16S and fungal ITS regions were processed in an Illumina MiSeq sequencer (run center: KYOTO-HE; 15% PhiX spike-in). Because the quality of forward sequences is generally higher than that of reverse sequences in Illumina sequencing, we optimized the MiSeq run setting in order to use only forward sequences. Specifically, the run length was set 271 forward (R1) and 31 reverse (R4) cycles in order to enhance forward sequencing data: the reverse sequences were used only for discriminating between 16S and ITS1 sequences based on the sequences of primer positions.

### Bioinformatics

The raw sequencing data were converted into FASTQ files using the program bcl2fastq 1.8.4 distributed by Illumina. The output FASTQ files were demultiplexed with the program Claident v0.2.2017.05.22 (Tanabe & Toju 2013; Tanabe 2017), by which sequencing reads whose 8-mer index positions included nucleotides with low (< 30) quality scores were removed. The sequencing data were deposited to DNA Data Bank of Japan (DDBJ) (DRA006845). Only forward sequences were used in the following analyses after removing low-quality 3’-ends using Claident. Noisy reads (Tanabe 2017) were subsequently discarded and then denoised dataset consisting of 2,041,573 16S and 1,325,199 ITS1 reads were obtained.

For each dataset of 16S and ITS1 regions, filtered reads were clustered with a cut-off sequencing similarity of 97% using the program VSEARCH (Rognes et al. 2014) as implemented in Claident. The operational taxonomic units (OTUs) representing less than 10 sequencing reads were subsequently discarded. The molecular identification of the remaining OTUs was performed based on the combination of the query-centric auto-*k*-nearest neighbor (QCauto) method (Tanabe & Toju 2013) and the lowest common ancestor (LCA) algorithm (Huson et al. 2007) as implemented in Claident. Note that taxonomic identification results based on the combination of the QCauto search and the LCA taxonomic assignment are comparable to, or sometimes more accurate than, those with the alternative approaches (Tanabe & Toju 2013; Toju et al. 2016b; Toju et al. 2016c). In total, 5,351 prokatyote (bacterial or archaeal) OTUs and 1,039 fungal OTUs were obtained for the 16S and ITS1 regions, respectively (Supplementary Data 1). The UNIX codes used in the above bioinformatic pipeline are available as Supplementary Data 2.

For each combination of target region (16S or ITS1) and sample type (root or soil), we obtained a sample × OTU matrix, in which a cell entry depicted the number of sequencing reads of an OTU in a sample (Supplementary Data 3). The cell entries whose read counts represented less than 0.1% of the total read count of each sample were removed to minimize effects of PCR/sequencing errors (Peay et al. 2015). The filtered matrix was then rarefied to 1,000 reads per sample using the “rrarefy” function of the vegan 2.4-1 package (Oksanen et al. 2012) of R 3.4.3 (R-Core-Team 2017). Samples with less than 1,000 reads were discarded in this process: the numbers of samples in the rarefied sample × OTU matrices were 119, 128, 117, and 128 for root prokaryote, root fungal, soil prokaryote, and soil fungal matrices, respectively (Supplementary Data 4).

### Prokaryote and Fungal Community Structure

Relationship between the number of sequencing reads and that of detected OTUs was examined for each dataset (root prokaryote, root fungal, soil prokaryote, or soil fungal dataset) with the “rarecurve” function of the R vegan package. Likewise, relationship between the number of samples and that of OTUs was examined with the vegan “specaccum” function. For each dataset, difference in OTU compositions among “green”, “yellow”, and “no leaf” soybean individuals was examined by the permutational analysis of variance [PERMANOVA; Anderson (2001)] with the vegan “adonis” function (10,000 permutations). To control effects of sampling positions (lines) on the community structure, the information of sampling sets (set 1 or set 2) was included as an explanatory variable in the PERMANOVA. The variation in OTU compositions was visualized with nonmetric multidimensional scaling (NMDS) using the vegan “metaMDS” function. To examine potential relationship between root/soil microbial community structure and plant biomass, an additional PERMANOVA was performed for each dataset. The information of sampling sets was included in the models. To explore signs of spatial autocorrelation in the community data, a Mantel’s correlogram analysis was performed with the vegan “mantel.correlog” function. The “Bray-Curtis” metric of *β*-diversity was used in the PERMANOVA, NMDS, and Mantel’s correlogram analyses.

### Screening of Host-state-specific OTUs

To explore prokaryote/fungal OTUs that preferentially occurred on/around “green”, “yellow”, or “no leaf” soybean individuals, a randomization test was performed by shuffling the plant state labels in each of the root prokaryote, root fungal, soil prokaryote, and soil fungal data matrices (100,000 permutations). We then evaluated preference of a prokaryote/fungal OUT (*i*) for a plant state (*j*) (“green”, “yellow”, or “no leaf”) as follows:

*Preference* (*i, j*) = [*N*_observed_ (*i, j*) – Mean (*N*_ranodomized_ (*i, j*))] / SD (*N*_ranodomized_ (*i, j*)), where *N*_observed_ (*i, j*) denoted the mean number of the sequencing reads of OTU *i* among state *j* soybean samples in the original data, and the Mean (*N*_ranodomized_ (*i, j*)) and SD (*N*_ranodomized_ (*i, j*)) were the mean and standard deviation of the number of sequencing reads for the focal OTU– plant state combination across randomized matrices. Regarding this standardized preference index, values larger than three generally represent strong preferences [false discovery rate (FDR) < 0.05; Toju et al. (2016b)] (Supplementary Fig. 5): hence, we listed OTUs whose preference values exceeded three.

### Microbe–microbe Networks

To examine how prokaryote and fungal OTUs co-occurred in root or soil samples, a co-abundance network analysis was performed based on the sparse inverse covariance estimation for ecological association inference (Spiec-Easi) method (Kurtz et al. 2015). In each of the root and soil sample analyses, the input data matrix was prepared by merging the sample × OTU matrices of prokaryotes and fungi. As inferences of co-abundance patterns were unavailable for rare OTUs, only the OTUs detected from 30 or more samples were retained in the input matrices. For each of the root and soil data matrices, a co-abundance analysis was performed with the “spiec.easi” function of the R “SpiecEasi” package (Kurtz et al. 2015). The networks depicting the co-abundance patterns were drawn using the R “igraph” package (Csardi & Nepusz 2006).

## RESULTS

### Prokaryotes and Fungal Community Structure

On average, 107.9 (SD = 18.0), 25.4 (SD = 8.9), 172.5 (SD = 17.3), and 78.3 (SD = 10.5) OTUs per sample were observed, respectively, from the root prokaryote, root fungal, soil prokaryote, and soil fungal dataset after filtering and rarefaction steps (Supplementary Fig. 2). The total number of OTUs observed was 1387, 346, 1191, and 769 for the root prokaryote, root fungal, soil prokaryote, and soil fungal datasets, respectively (Supplementary Fig. 3).

**FIGURE 2.**
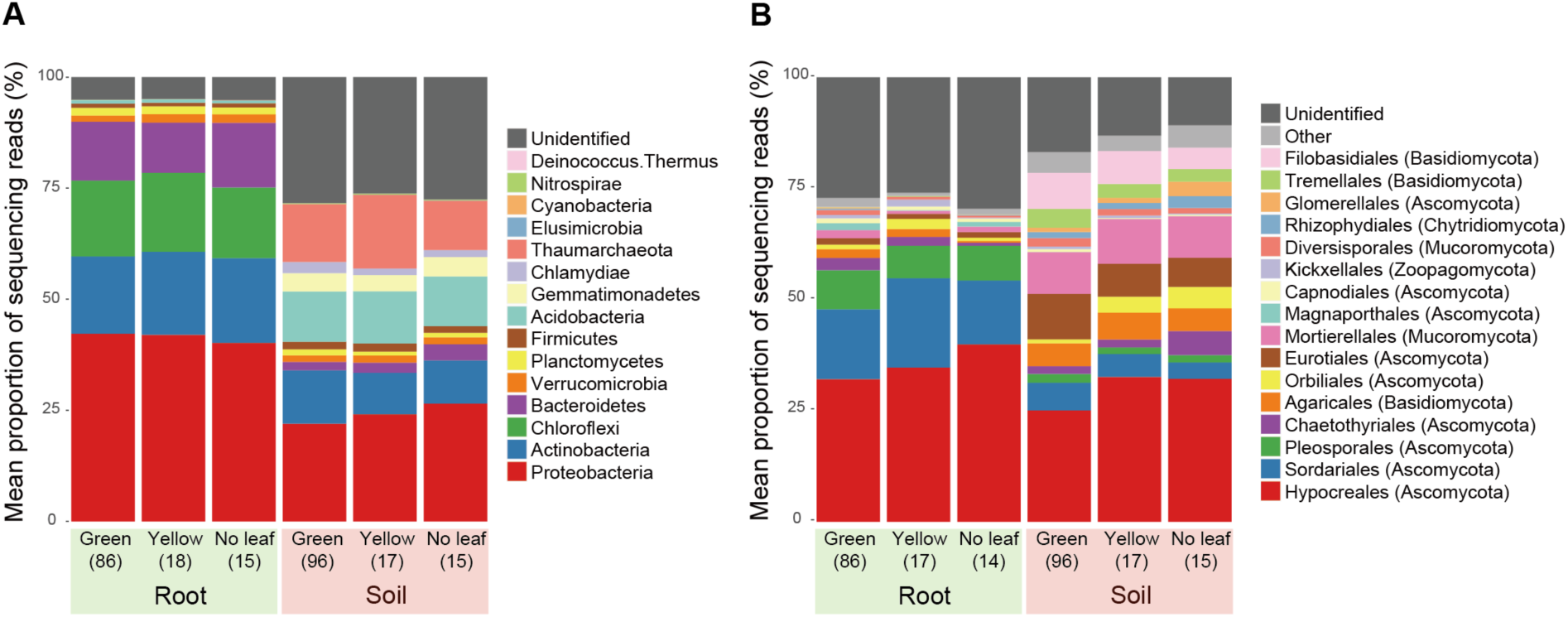
Prokaryote and fungal community structure. **(A)** Phylum-level compositions of prokaryotes in the root and soil datasets. Mean proportions of sequencing reads are shown for each taxa. The numbers of the samples from which sequencing data were successfully obtained are shown in the parentheses. **(B)** Order-level compositions of fungi in the root and soil datasets.

**FIGURE 3.**
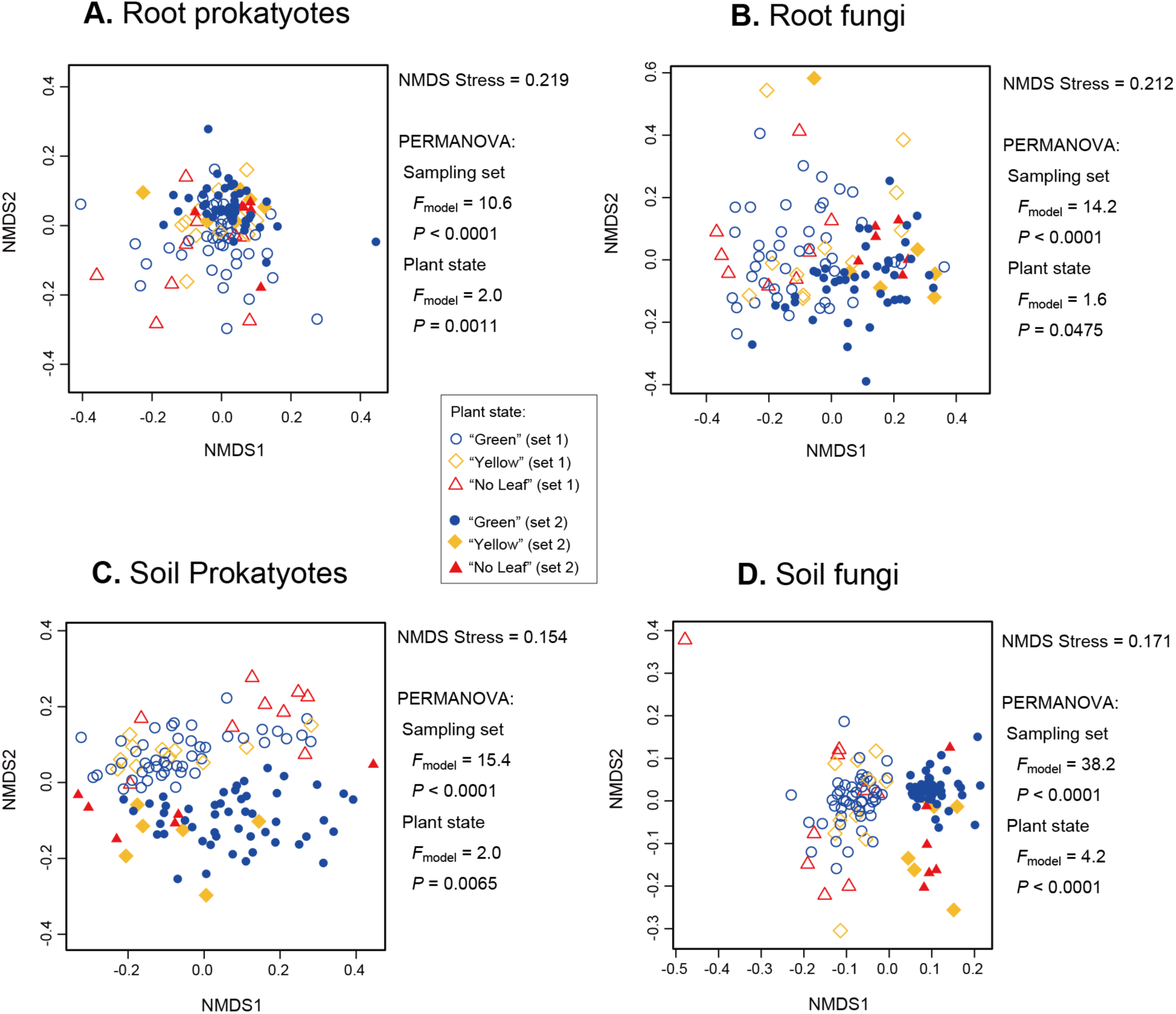
Diversity of microbiome structures among samples. **(A)** NMDS of the root prokaryote dataset. The results of the PERMANOVA, in which sampling set (“set 1” or “set 2”) and plant state (“green”, “yellow”, or “no leaf”) were included as explanatory variables, are shown. **(B)** NMDS of the root fungal dataset. **(C)** NMDS of the soil prokaryote dataset. **(D)** NMDS of the soil fungal dataset.

In the soybean field, the prokaryote community on roots was dominated by the bacterial classes Proteobacteria, Actinobacteria, Chloroflexi, and Bacteroidetes, while that of rhizosphere soil consisted mainly of Proteobacteria, Actinobacteria, and Acidobacteria, and the archaeal lineage Thaumarchaeota (Fig. 2A). The fungal community of roots was dominated by the fungal orders Hypocreales, Sordariales, Plesporales, while that of soil consisted mainly of Hypocreales, Agaricales, Eurotiales, Mortierellales, and Filobasidiales (Fig. 2B). Regarding the order level compositions of fungi in the rhizosphere soil, the proportion of Orbiliales reads was much higher in “yellow” (3.62 %) and “no leaf” (4.82 %) soybean individuals than in “green” ones (0.89 %) (Fig. 2).

In each dataset (i.e., root prokaryote, root fungal, soil prokaryote, or soil fungal data), microbial community structure varied among “green”, “yellow”, or “no leaf” soybean individuals, although the effects of sampling sets on the community structure were much stronger (Fig. 3). Even within each sampling set, spatial autocorrelations of bacterial/fungal community structure were observed (Supplementary Fig. 4). Significant relationships between microbial community structure and soybean biomass were observed in the soil prokaryote and soil fungal datasets but not in the root prokaryote and root fungal datasets (Table 1).

**TABLE 1.** Relationship between prokaryote/fungal community structure and the biomass of soybean individuals.

| Variable | df | $F_{\text{model}}$ | $P$ |
| --- | --- | --- | --- |
| Root prokaryotes |  |  |  |
| Sampling set | 1 | 10.4 | < 0.0001 |
| Dry mass | 1 | 1.3 | 0.1139 |
| Root fungi |  |  |  |
| Sampling set | 1 | 14.0 | < 0.0001 |
| Dry mass | 1 | 0.6 | 0.8267 |
| Soil prokaryotes |  |  |  |
| Sampling set | 1 | 15.4 | < 0.0001 |
| Dry mass | 1 | 3.1 | 0.002 |
| Soil fungi |  |  |  |
| Sampling set | 1 | 36.7 | < 0.0001 |
| Dry mass | 1 | 2.2 | 0.0145 |

For each dataset (i.e., root prokaryote, root fungal, soil prokaryote, or soil fungal data), a PEMANOVA model of community structure was constructed. The information of the sampling set (“set 1” or “set 2”) and the dry mass of host soybean individuals were included as explanatory variables.

**FIGURE 4.**
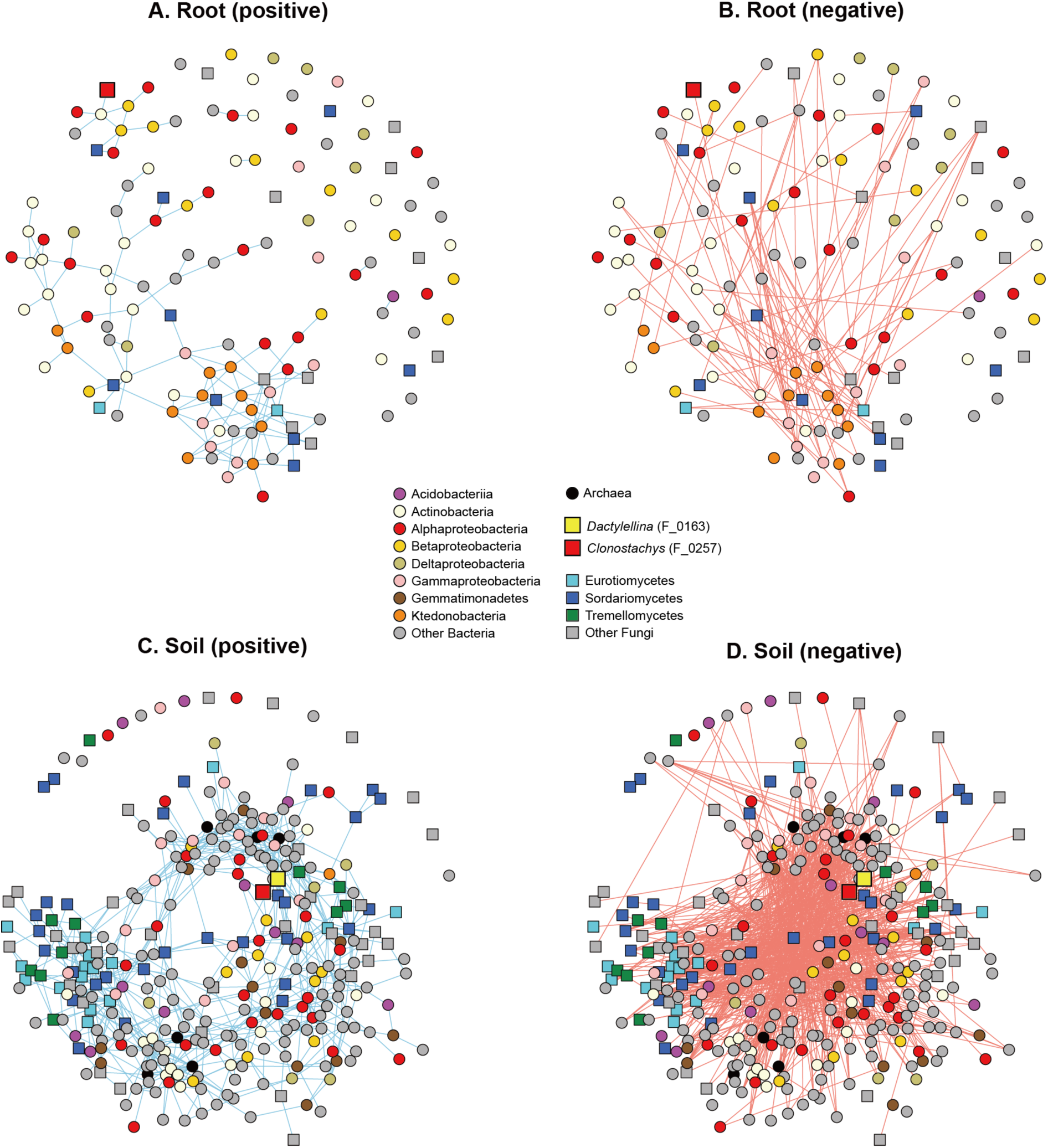
Microbe–microbe co-abundance networks. **(A)** Positive co-abundance network of the root microbiome data. A pairs of OTUs linked by a blue line frequently co-occurred in the same soybean samples. **(B)** Negative co-abundance network of the root microbiome data. A pairs of OTUs linked by a red line rarely co-occurred in the same soybean samples. **(C)** Positive co-abundance network of the soil microbiome data. **(D)** Negative co-abundance network of the soil microbiome data.

### Screening of Host-state-specific OTUs

In the root microbiome, only an unidentified fungal OTU showed a strong preference for “green” soybean individuals, while 18 bacterial and 4 fungal OTUs occurred preferentially on “no leaf” host individuals (Table 2). The list of the bacteria showing preferences for “no leaf” soybean individuals included OTUs whose 16S rRNA sequences were allied to those of *Dyella, Herbaspirillum, Labrys, Phenylobacterium, Gemmata, Chitinophaga, Pedobacter, Niastella*, and *Streptomyces* (Table 2). The four fungal OTUs showing preferences for “no leaf” hosts were unidentified basidiomycetes (Table 2).

**TABLE 2.**
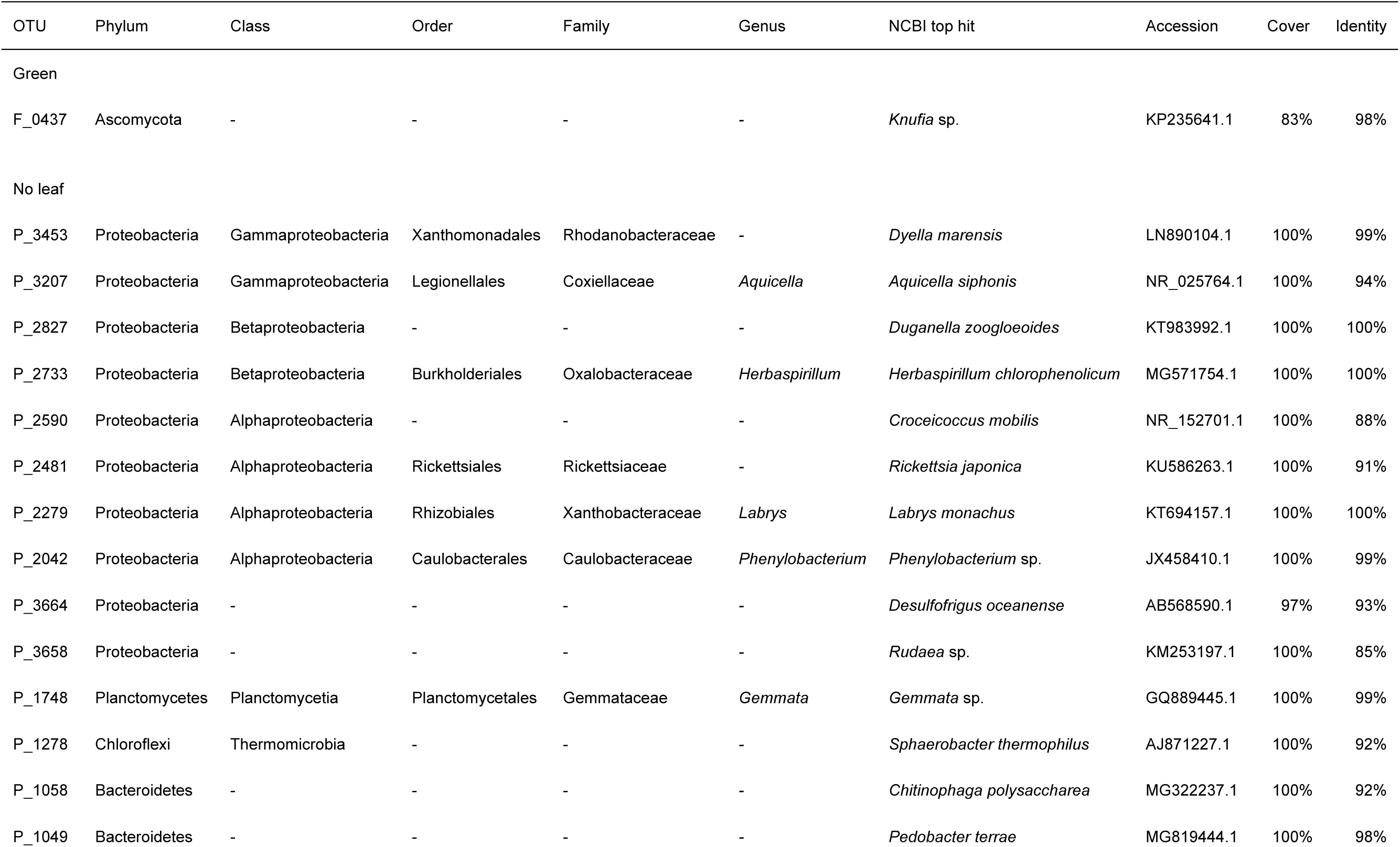
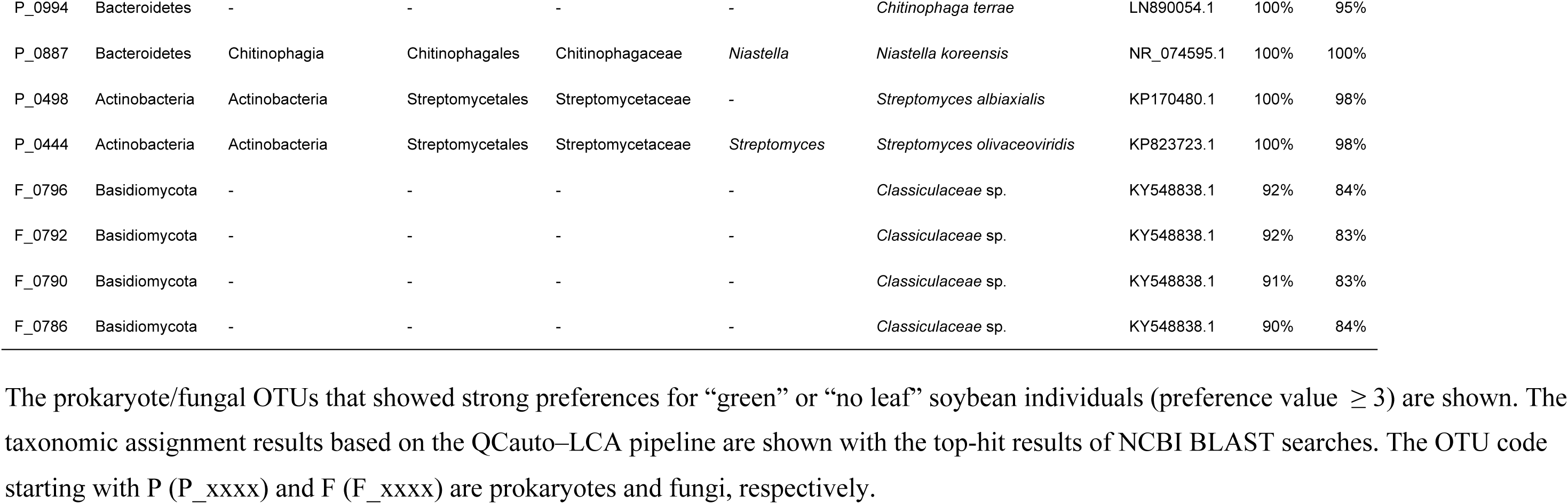
Prokaryote and fungal OTUs showing strong preferences for host states in the root microbiome datasets.

In the rhizosphere soil microbiome, seven prokaryote OTUs, including those belonging to Chloroflexi (e.g., *Sphaerobacteraceae* sp.) and Proteobacteria (*Kofleriaceae* sp.), occurred preferentially on “green” host individuals (Table 3). Likewise, five fungal OTUs, including those allied to basidiomycete yeasts in the genera *Solicoccozyma* and *Saitozyma*, showed preferences for “green” soybean individuals (Table 3). Results also revealed that 26 bacterial and 11 fungal OTUs had biased distributions in the rhizosphere of “no leaf” soybean individuals (Table 3). The list of microbes showing preferences for “no leaf” hosts included OTUs allied to bacteria in the genera *Pesudomonas, Nevskia, Cellvibrio, Massilia, Duganella, Novosphingobium, Mucilaginibacter*, and *Flavobacterium* and OTUs allied to fungi in the genera *Burgoa, Clonostachys, Plectosphaerella, Xylaria, Dactylellina, Talaromyces, Cladosporium, Alternaria*, and *Peniophora* (Table 3). The list of microbes that preferentially occurred on “no leaf” hosts involved OTUs with high sequence similarity to the nematophagous fungi, *Clonostachys rosea* (Hypocreales) and *Dactylellina* sp. (Orbiliales) (Table 3). The reads of the *Clonostachys* (F_0257) and *Dactylellina* (F_0163) OTUs, respectively, represented 9.5% and 3.5% of the sequencing reads of “no leaf” samples (Supplementary Data 5). The indices of preferences for “yellow” soybean individuals are shown in Supplementary Data 5.

**TABLE 3.**
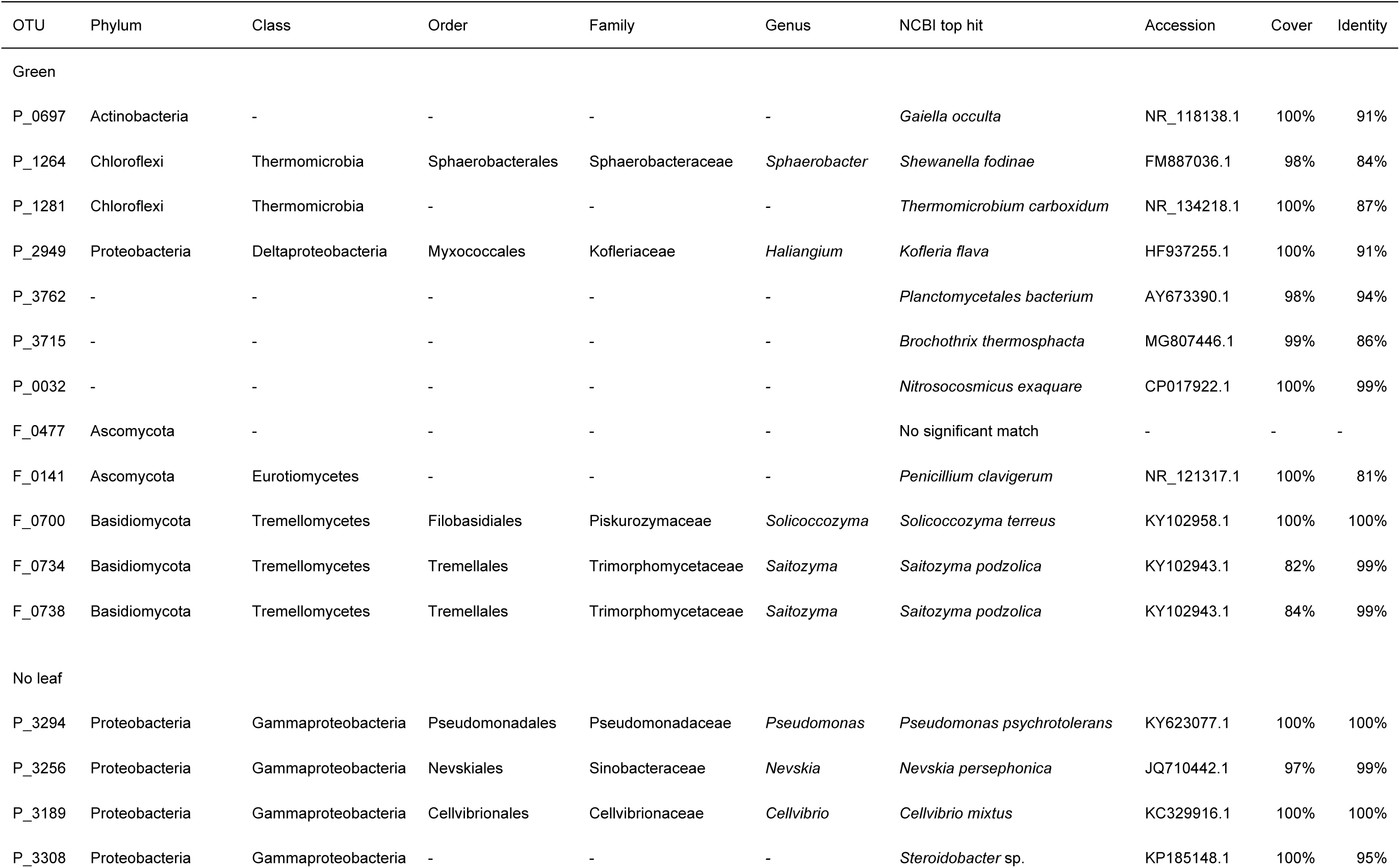
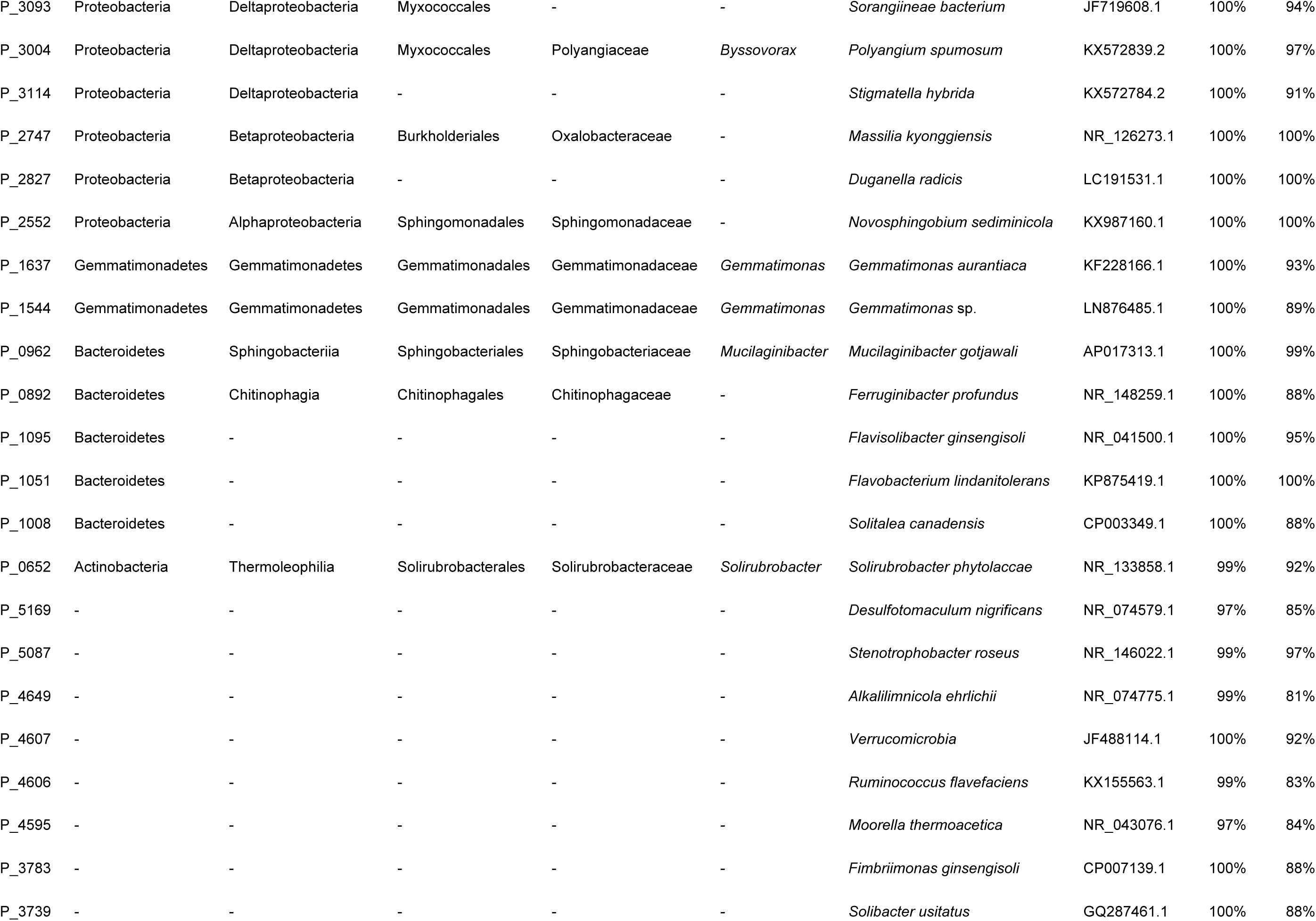
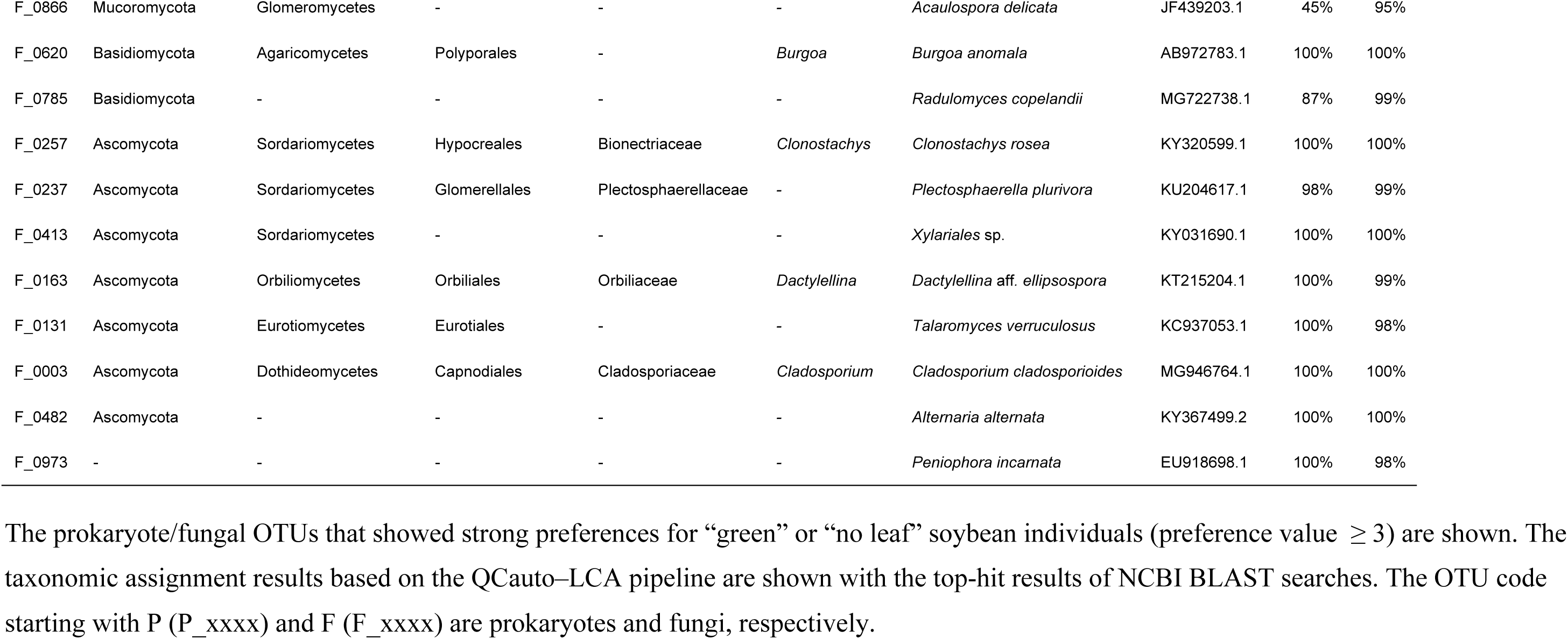
Prokaryote and fungal OTUs showing strong preferences for host states in the soil microbiome datasets.

### Microbe–microbe Networks

The structure of microbe–microbe networks (Fig. 4) were more complicated in the soil microbiome data (Fig. 4C-D) than in the root microbiome data (Fig. 4A-B). Within the network representing co-abundance of microbes across root samples, the *Clonostachys* OTU (F_0257) had a significant link with a *Streptomyces* OTU, while *Dactylellina* was absent from the root microbiome network data (Fig. 4A). Within the positive co-abundance network of the rhizosphere soil microbiome (Fig. 4C), the *Clonostachys* (F_0257) and *Dactylellina* (F_0163) nematophagous fungal OTUs were connected with each other (Table 4). In addition, the *Clonostachys* OTU was linked with two bacterial OTUs (*Ralstonia* and Rhizobiales) and fungal OTUs in the genera *Calonectria* and *Purpureocillium* (Table 4). Likewise, the *Dactylellina* OTU was connected also with two Alphaproteobacterial OTUs and a bacterial OTU allied to *Nitrospira japonica* as well as fungal OTUs in the genera *Rhizophydium, Pochonia, Purpureocillium* (Table 4).

**TABLE 4.**
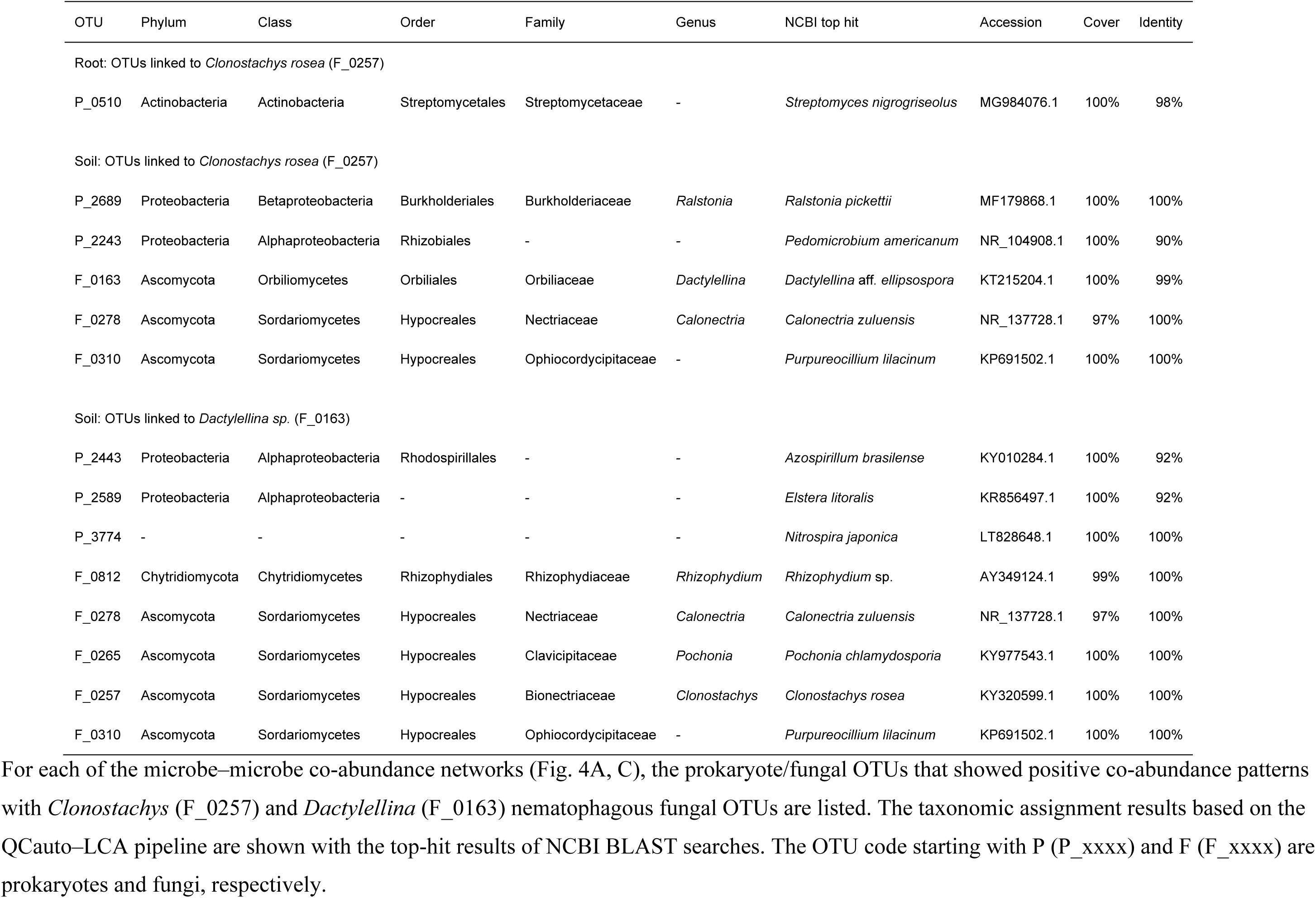
Prokaryote/fungal OTUs linked to nematophagous fungi in the microbe–microbe networks.

| OTU | Phylum | Class | Order | Family | Genus | NCBI top hit | Accession | Cover | Identity |
| --- | --- | --- | --- | --- | --- | --- | --- | --- | --- |
| Root: OTUs linked to *Clonostachys rosea* (F_0257) |  |  |  |  |  |  |  |  |  |
| P_0510 | Actinobacteria | Actinobacteria | Streptomycetales | Streptomycetaceae | - | *Streptomyces nigrogriseolus* | MG984076.1 | 100% | 98% |
| Soil: OTUs linked to *Clonostachys rosea* (F_0257) |  |  |  |  |  |  |  |  |  |
| P_2689 | Proteobacteria | Betaproteobacteria | Burkholderiales | Burkholderiaceae | *Ralstonia* | *Ralstonia pickettii* | MF179868.1 | 100% | 100% |
| P_2243 | Proteobacteria | Alphaproteobacteria | Rhizobiales | - | - | *Pedomicrobium americanum* | NR_104908.1 | 100% | 90% |
| F_0163 | Ascomycota | Orbiliomycetes | Orbiliales | Orbiliaceae | *Dactylellina* | *Dactylellina* aff. *ellipsospora* | KT215204.1 | 100% | 99% |
| F_0278 | Ascomycota | Sordariomycetes | Hypocreales | Nectriaceae | *Calonectria* | *Calonectria zuluensis* | NR_137728.1 | 97% | 100% |
| F_0310 | Ascomycota | Sordariomycetes | Hypocreales | Ophiocordycipitaceae | - | *Purpureocillium lilacinum* | KP691502.1 | 100% | 100% |
| Soil: OTUs linked to *Dactylellina* sp. (F_0163) |  |  |  |  |  |  |  |  |  |
| P_2443 | Proteobacteria | Alphaproteobacteria | Rhodospirillales | - | - | *Azospirillum brasilense* | KY010284.1 | 100% | 92% |
| P_2589 | Proteobacteria | Alphaproteobacteria | - | - | - | *Elstera litoralis* | KR856497.1 | 100% | 92% |
| P_3774 | - | - | - | - | - | *Nitrospira japonica* | LT828648.1 | 100% | 100% |
| F_0812 | Chytridiomycota | Chytridiomycetes | Rhizophydiales | Rhizophydiaceae | *Rhizophydium* | *Rhizophydium* sp. | AY349124.1 | 99% | 100% |
| F_0278 | Ascomycota | Sordariomycetes | Hypocreales | Nectriaceae | *Calonectria* | *Calonectria zuluensis* | NR_137728.1 | 97% | 100% |
| F_0265 | Ascomycota | Sordariomycetes | Hypocreales | Clavicipitaceae | *Pochonia* | *Pochonia chlamydosporia* | KY977543.1 | 100% | 100% |
| F_0257 | Ascomycota | Sordariomycetes | Hypocreales | Bionectriaceae | *Clonostachys* | *Clonostachys rosea* | KY320599.1 | 100% | 100% |
| F_0310 | Ascomycota | Sordariomycetes | Hypocreales | Ophiocordycipitaceae | - | *Purpureocillium lilacinum* | KP691502.1 | 100% | 100% |

## DISCUSSION

Based on Illumina sequencing, we herein compared root-associated/rhizosphere microbial communities between soybean individuals infected by root-knot nematodes and those showing no symptoms. The results indicated that, in both soybean roots and rhizosphere soil, prokaryote and fungal community structures significantly varied depending on host plant states (Figs. 2 and 3). We further performed statistical analyses for screening prokaryote and fungal OTUs preferentially associated with infected and benign soybean host individuals (Tables 2-3; Fig. 4). The results are based on purely descriptive data and hence they, in principle, are not direct evidences of interactions among plants, nematodes, and microbiomes. Moreover, as this study provided only “snap-shot” information of microbiome structure at the end of a growing season, further studies uncovering temporal microbiome dynamics throughout the growing season of soybeans are awaited. Nonetheless, as detailed below, the statistical analyses suggest assembly of diverse anti-nematode bacteria and fungi from indigenous microbial communities in the soybean field, providing a basis for exploring ways to reduce damage by root-knot nematodes with those indigenous functional microbes.

Within the root microbiome analyzed, various taxonomic groups of bacteria preferentially occurred on “no leaf” soybean samples (Table 2). Among them, the genus *Streptomyces* is known to involve some species that suppress nematode populations, potentially used as biological control agents for root-knot nematodes (Esnard et al. 1995; Chubachi et al. 1999; Siddiqui & Mahmood 1999; Samac & Kinkel 2001). In contrast, *Herbaspirillum, Rickettsia, Chitinophaga*, and *Pedobacter* have been reported as symbionts of nematodes, potentially playing beneficial roles for host nematodes (Tian et al. 2011; Baquiran et al. 2013; Cheng et al. 2013). Thus, results of these statistical analyses should be interpreted with caution, as they are likely to highlight not only prospective microbe potentially parasitizing on pests/pathonges but also microbes that can form mutualistic interactions with disease agents.

Within the soybean rhizosphere soil microbiome, diverse taxonomic groups of not only bacteria but also fungi preferentially occurred around “no leaf” soybean individuals (Table 3). Among them, *Pseudomonas* has been known to suppress root-knot nematode populations (Siddiqui & Ehteshamul-Haque 2001; Siddiqui & Shaukat 2004) potentially by producing hydrogen cyanide (Siddiqui et al. 2006) or extracellular protease (Siddiqui et al. 2005), but interactions with root-knot nematodes have not yet been examined for other bacteria preferentially found in the rhizosphere of “no leaf” soybean individuals. Meanwhile, the list of the fungal OTUs frequently observed in the rhizosphere of “no leaf” soybeans included some fungi whose ability to suppressing nematode populations had been well documented (Table 3). *Clonostachys rosea*, for example, has been known as a prospective biological control agent of plant-and animal-pathogenic nematodes (Zou et al. 2010b; Baloyi et al. 2011). An observational study based on green fluorescent protein imaging has indicated that the conidia of the fungus adhere to nematode cuticle and their germ tubes penetrate nematode bodies, eventually killing the invertebrate hosts (Zhang et al. 2008). The fungus is also known to produce a subtilisin-like extracellular protease, which play an important role during the penetration of nematode cuticles (Zou et al. 2010a). In addition to *Clonostachys*, our analysis highlighted another nematophagous fungus in the genus *Dactylellina* (teleomorph = *Orbilia*). Species in the genus and many other fungi in the order Orbiliales produce characteristic trap structures with their hyphae to prey on nematodes (Liu et al. 2005; Yang et al. 2007b; Yu et al. 2012), often nominated as prospective biological control agents (Yang et al. 2007a; Timper 2011; Moosavi & Zare 2012).

An additional analysis focusing on *Clonostachys* and *Dactylellina* highlighted bacteria and fungi that frequently co-occur with the nematophagous fungi (Fig. 4). In the root microbiome, *Clonostachys* and a *Streptomyces* OTU showed positively correlated distributions across soybean samples (Table 4). In the rhizosphere microbiome, *Clonostachys* and *Dactylellina* showed significant co-abundance patterns (Table 4). Moreover, in the soil, the two nematophagous fungi co-occurred frequently with other taxonomic groups of nematophagous fungi such as *Purpureocillium, Pochonia*, and *Rhizophydium* (Table 4; Fig. 5). Among them, fungi in the genus *Purpureocillium* (Hypocreales: Ophiocordycipitaceae) have been known to suppress plant parasitic nematodes, insect pests, and oomycete phytopathogens (Lopez-Lima et al. 2014; Goffré & Folgarait 2015; Lopez & Sword 2015; Wang et al. 2016) and their genome sequences have been analyzed for understanding the physiological mechanisms of the pest/pathogen suppression (Prasad et al. 2015; Wang et al. 2016; Xie et al. 2016). As one of *Purpureocillium* species (*P. liacinum*) can form symbiotic interactions with plants as endophytes (Lopez et al. 2014; Lopez & Sword 2015), it has been recognized as promising biological control agents for commercial use (Wang et al. 2016). Another Hypocreales genus, *Pochonia* (previously placed in the genus *Verticillium*; teleomorph = *Metacordyceps*; Clavicipitaceae) has been known as nematophagous as well and they can kill eggs and females of root-knot (*Meloidogyne* spp.) and cyst (*Globodera* spp.) nematodes (Atkins et al. 2003; Tobin et al. 2008; Niu et al. 2009; Siddiqui et al. 2009). *Pochonia* fungi, especially *P. chlamydosporia*, are also endophytic and hence they have been used in agriculture (Maciá-Vicente et al. 2009a; Maciá-Vicente et al. 2009b; Escudero & Lopez-Llorca 2012; Larriba et al. 2014). Species in the chytrid genus *Rhizophydium* involve species that utilize nematodes as parasites or saprophytes (Barr 1970; Esser 1983). They are known to explore host nematodes in the form of zoospores (Esser 1983). Overall, our results suggest that indigenous anti-nematode or nematophagous microbes can form consortia in soil ecosystems of soybean fields. It is important to note that the members of the consortia do not necessarily interact with each other directly: i.e., they may merely share habitat preferences (Dickie 2007; Peay et al. 2015; Toju et al. 2016c). However, the inferred structure of microbe–microbe networks helps us understand overall consequences of ecological processes in microbiomes (Toju et al. 2018).

**FIGURE 5.**
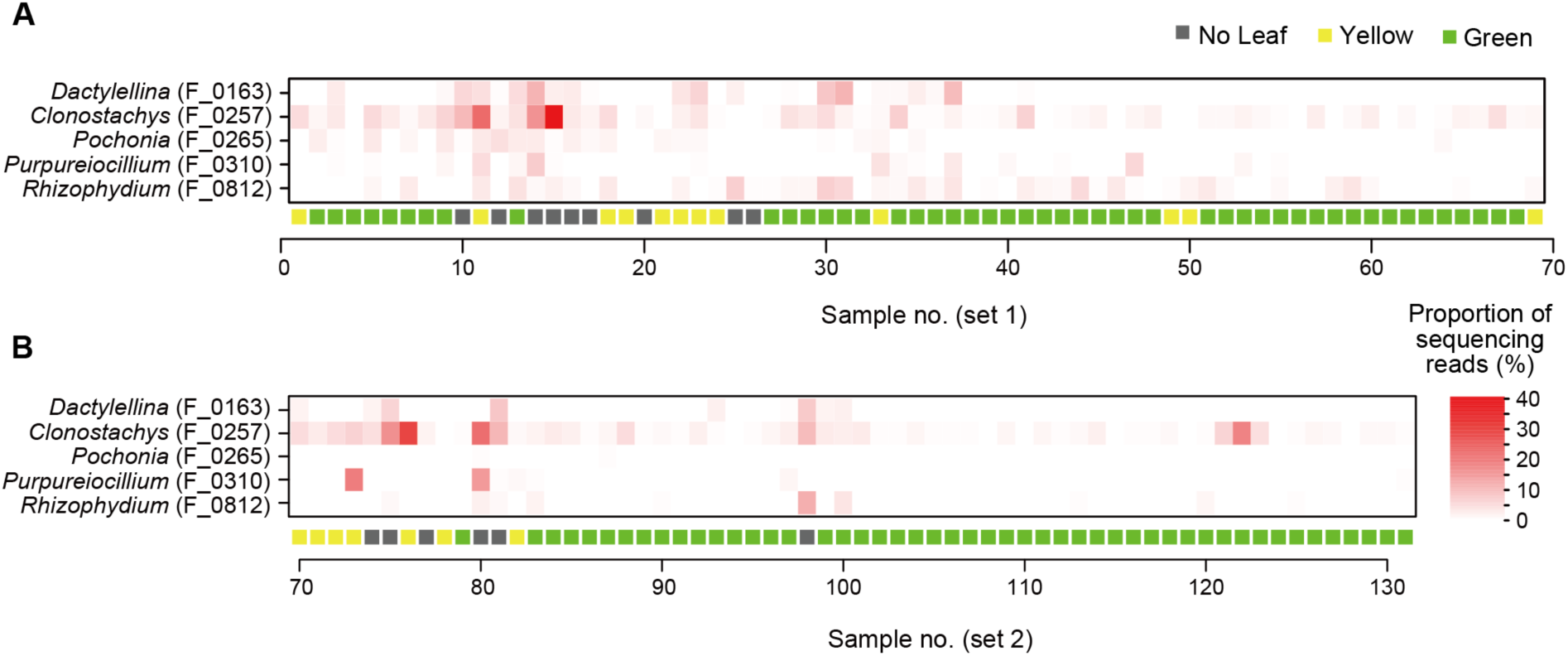
Spatial distribution of nematophagous fungal OTUs. **(A)** Sampling set 1. For 39 each soybean individual, the proportions of sequencing reads representing nematophagous 40 fungal OTUs are shown. **(B)** Sampling set 2.

Along with the consortia of anti-nematode microbes, an OTU in the genus *Calonetria*, which causes leaf blight, wilt, and root rot of various plant species (Kuruppu et al. 2004; Vitale et al. 2013), was frequently observed (Table 4). The phytopathogenic fungus might have attacked soybean individuals weakened by root-knot nematodes. Alternatively, *Calonectria* may have infected host soybeans earlier than root-knot nematodes, followed by the emergence of nematodes and their exploiters (i.e., anti-nematode microbes). Given that fungi can interact with each other both antagonistically and mutualistically in the soil (Verma et al. 2007; Toju et al. 2016a), direct interactions between *Calonectria* and nematophagous fungi in the genera *Clonostachys, Dactylellina, Purpureocillium, Pochonia*, and *Rhizophydium* are of particular interest. Studies examining potential interactions involving soybeans, root-knot nematodes, anti-nematode bacteria/fungi, and *Calonectria* will help us understand ecological processes that structure consortia of nematophagous fungi.

Although this study did not evaluate potential effects of background environmental conditions (e.g., soil pH and inorganic nitrogen concentration) on microbiome structure, management of edaphic conditions are expected to have great impacts on dynamics of anti-nematode microbiomes. A number of studies have explored ways to suppress nematode populations by optimizing cropping systems (Barker & Koenning 1998). Crop rotation, in which planting of a crop variety and that of nematode-resistant varieties/species are rotated, has been recognized as an effective technique for regulating root-knot and cyst nematode populations (Nusbaum & Ferris 1973; Koenning et al. 1993; Chen 2007). In contrast, long-term continual cropping in soybean monoculture fields can increase anti-nematode bacteria and fungi (e.g., *Pseudomonas, Purpureocillium*, and *Pochonia*), potentially resulting in lowered densities of cyst nematodes (Hamid et al. 2017). Tillage regimes (Thomas 1978; Okada & Harada 2007; Donald et al. 2009) and introduction of organic matter (e.g., alfalfa leaves or crop residue) (Hershman & Bachi 1995; Jaffee 2004, 2006) have great impacts on nematode densities in farmlands, but their effects vary considerably among studies (Barker & Koenning 1998). In addition, because plant individuals infected by nematodes can show highly aggregated distributions at a small spatial scale within a farmland (Fig. 1D), tillage can promote the spread of plant damaging nematodes (Gavassoni et al. 2001). Frequent tillage may have negative impacts on populations of nematophagous fungi as a consequence of hyphal fragmentation [cf. Verbruggen & Toby Kiers (2010)], but such destructive effects on fungal communities have not yet been tested intensively. Given that microbiome structures were not taken into account in most previous studies evaluating effects of cropping systems on nematode suppression [but see Hamid et al. (2017)], more insights into relationship between agroecosystem management and indigenous (native) microbiome dynamics are required for building reproducible ways for developing disease-suppressive soil.

We herein found that consortia of anti-nematode bacteria and fungi could develop at a small spatial scale within a field of soybeans infected by root-knot nematodes. Taking into account the diversity of those anti-nematode microbes observed in this study, multiple biological control agents are potentially available *in situ* without introducing exogenous ones depending on base compositions and conditions of indigenous microbiomes within and around a focal farmland. In this respect, design of cropping systems (e.g., crop rotations, tillage frequencies, and inputs of fertilizer or organic matter) is of particular importance in activating and maximizing ecosystem functions that stem from resident microbial diversity (Toju et al. 2018). Because those indigenous microbes, in general, have adapted to local biotic and abiotic environments, their populations are expected to persist more stably than exogenous microbes artificially introduced to a target agroecosystem [see Mendes et al. (2013) for reviews of the success/failure of microbial introduction]. Elucidating relationship between cropping systems and microbiome processes is the key to design disease-suppressive agroecosystems.

## AUTHOR CONTRIBUTIONS

HT designed the work. HT and YT performed fieldwork. HT conducted molecular experiment and analyzed the data. HT wrote the manuscript with YT.

## ACKNOWLEDGEMENTS

We thank Tatsuhiko Shiraiwa for his support in fieldwork and Mizuki Shinoda, Ko Mizushima, Sarasa Amma, and Hiroki Kawai for their support in molecular experiments. We are also grateful to Ryoji Shinya for his advice on the biology of nematodes. This work was financially supported by JSPS KAKENHI Grant (15KT0032) and JST PRESTO (JPMJPR16Q6) to HT.

## SUPPLEMENTARY MATERIAL

The Supplementary Material for this article can be found online at: XXXX.

